# Physiological, transcriptomic and metabolomic responses of the marine diatom *Phaeodactylum tricornutum* to auxin

**DOI:** 10.1101/2024.11.24.623293

**Authors:** Yu-Ting Chen, Dong-Sheng Zhao, Xiao-Li Liu, Huan Yang, Run-Ze Gu, Nan Li, Xiufeng Yan, Hui-Xi Zou

## Abstract

Auxin, an essential phytohormone, is widely used to promote plant growth and development. However, the effect of auxin on diatoms and its mechanism remain underexplored. Here, we studied the impact of auxin 3-indoleacetic acid (IAA) on the marine diatom *Phaeodactylum tricornutum* and the underlying molecular mechanisms. We found that 5 μg L^-1^ of IAA promotes the growth of *P. tricornutum* in a time-dependent manner. Treatment with IAA resulted in significant changes in photosynthetic pigments and malondialdehyde, chlorophyll fluorescence, and antioxidant enzyme activities. In addition, transcriptomics showed that IAA exposure leads to upregulation of a large number of differential genes (DEGs) in carbon fixation and porphyrin metabolism, several of which were verified by qPCR. Furthermore, corresponding metabolites of these pathways were also validated by metabolomic analysis. Thus, IAA exhibits growth-promotion effect on the diatom mainly through increasing photosynthetic carbon sequestration and the expression of genes in porphyrin synthesis. Our results provide key data on the action mechanism of IAA in promoting the growth of a diatom.

## 1. INTRODUCTION

Diatoms are significant contributors to the aquatic food chain, supporting various organisms such as fish, copepods, and shellfish [1]. They are also promising alternatives for large-scale biodiesel production due to the accumulation of lipids and long-chain polyunsaturated fatty acids [2, 3]. Given the substantial commercial values, large-scale cultivation of diatoms has gained growing attention [4].

3-Indoleacetic acid (IAA), first discovered in maize [5], is the primary auxin in plants that regulates numerous development and growth events [6]. Auxin is evolutionary conserved as the enzymes responsible for its biosynthesis present in the common ancestor of land plants and Charophyte [7, 8]. In algae, the role of auxin had been reported in *Pleurochrysis portfolio* and *Chlorella sorokiniana*, demonstrating a positive correlation between auxin naphthalene acetate (NAA) and chlorophyll and biomass [9]. In the freshwater microalgae *Scenedesmus sp*., IAA and 2, 4-D promote growth by increasing chlorophyll content [10]. However, studies on the effects of auxin on diatoms and their mechanisms have lagged far behind.

The stimulation of microalgae growth by auxin represents a viable strategy for enhancing biomass and lipid production in microalgae [11]. The production of biofuels from microalgae has gained considerable attention as a viable alternative to fossil fuels. The marine diatom *Phaeodactylum tricornutum* plays a vital role in global ecosystems and is a promising source of energy[12]. However, the cost of large-scale microalgal production remains prohibitive. Therefore, improving the efficiency of microalgae production is the key for its commercial viability [13].

Here, we assessed the impact of IAA on the growth of *P. tricornutum* and found that 5 μg L^-1^ of IAA significantly enhances the various physiological parameters. Transcriptomics and metabolomics analyses were also conducted to elucidate the impacted pathways and differential metabolites, thereby outlining the role of IAA in fostering the growth of *P. tricornutum*. Our data will facilitate future research and pave the way for the utilization of *P. tricornutum* as a biofuel source.

## 2. MATERIALS AND METHODS

### 2.1. Chemicals

3-Indoleacetic acid (IAA) was purchased from Sigma-Aldrich. The stock solution was prepared using dimethyl sulfoxide (DMSO) and diluted to the specified concentrations .

### 2.2. Diatom strain and culture conditions

*Phaeodactylum tricornutum* Bohlin CCMP2561 was obtained from National Center for Marine Algae and Microbiota (USA). The diatom was grown at 20 ± 1 °C in f/2 medium under white fluorescent light (90 μmol m^-2^ s^-1^), 12 h:12 h dark-light cycle. All culture media were manually stirred two times a day.

### 2.3. Microalgal growth

The final concentration of DMSO in media was controlled at < 0.01% (v/v). IAA of a range of concentrations (100 pg L^-1^, 1 ng L^-1^, 10 ng L^-1^, 100 ng L^-1^, 5 μg L^-1^, 20 μg L^-1^, 40 μg L^-1^, and 100 μg L^-1^) was prepared in the f/2 medium in 12-well plates, to which 5 mL of *P. tricornutum* at the exponential phase was added. The cell density was set to 4 × 10^6^ cell mL^-1^. Cell density was calculated using measurement of optical density (OD) of 680 nm with the equation y = 100x + 1.12, where x refers to the OD_680_ and y refers to the cell density (10^6^ cells/mL) [14]. Samples were collected at time intervals of 24, 48, 72 and 96 h.

The growth promotion efficiency was calculated as described [15], using the following equation: Promotion = (N - N_0_) / N_0_ × 100%, where N and N_0_ are the cell density (cell mL^-1^) values in the treatment and control groups, respectively.

### 2.4. Pigment measurement

Ten ml cell culture was harvested by centrifugation at 5,000 rpm for 15 min. The pellet was re-suspended in 10 mL of 90% methanol, incubated at 55 °C for 15 min and again centrifuged for 15 min. The absorbance of the supernatant at 665, 652 and 470 nm was measured using a BioTek Epoch (BioTek, USA). The total chlorophyll and carotenoid concentration were calculated using the following equations [16]:

Chlorophyll *a* (mg L^-1^) = 16.82 × A_665_ - 9.28 × A_652_

Chlorophyll *b* (mg L^-1^) = 36.92 × A_652_ - 16.54 × A_665_

Carotenoids (mg L^-1^) = (100 × A_470_ - 1.91 × C_a_ - 95.15 × C_b_ ) / 225.

### 2.5. Chlorophyll fluorescence

Chlorophyll fluorescence including maximal photochemical efficiency (Fv/Fm), electron transfer rate (ETR), photochemical quenching (qP), and non-photochemical quenching (NPQ) were measured by a modulated chlorophyll fluorometer (Image- PAM, Heinz Walz GmbH, Effeltrich, Germany) [17]. Samples were collected at time intervals of 0, 24, 48, 72 and 96 h. After a 20 min dark period in ambient conditions, the chlorophyll fluorescence parameters were measured.

### 2.6. Oxygen consumption and production

Respiratory oxygen consumption and production were determined by a liquid oxygen electrode (Chlorolab 2, Hansatech, UK). The rates were expressed based on the mass of chlorophyll *a* in per unit volume (the initial biomass of Chl-*a* is 2 mg L^−1^). Samples were collected after 24, 48, 72 and 96 h, and 2 mL algal suspension was added into the reaction vessel (25 °C). The oxygen consumption was deducted when the net photosynthetic oxygen release was calculated.

### 2.7. Antioxidant enzyme activities and MDA content

The enzyme activity of superoxide dismutase (SOD), peroxidase (POD) and catalase (CAT), and the content of malondialdehyde (MDA) were determined by using commercial kits (Grace Biotechnology, Suzhou, China) according to the manufacturers’ instructions. Approximately 0.1 g algal samples (fresh weight, n = 3) were homogenized in liquid nitrogen and extracted with 1 mL of extract solution in the reagent kit. After centrifugation, the supernatant was used for the measurement.

### 2.8. Total RNA extraction and RNA-Seq

Total RNA was extracted by Trizol reagent according to the manufacture’s instruction (Invitrogen, Carlsbad, CA, USA). The RNA amount and purity of each sample were quantified using a NanoDrop ND-1000 (NanoDrop, Wilmington, USA). The integrity of the RNA was assessed using a Bioanalyzer 2100 (Agilent, CA, USA), which yielded RIN numbers greater than 7.0, and further confirmed through electrophoresis. Qualified RNA samples were submitted to Beijing Novogene Institute for library construction and Illumina sequencing. Following sequencing, reads of low and medium quality, containing junctions, and deficient in base information were eliminated. Subsequently, valid clean reads were extracted using the *P. tricornutum* CCAP 1055/1 reference genome obtained from the NCBI database. The reference genome index was constructed using Hisat2 v2.0.5, and the paired-end clean reads were compared with the reference genome. The obtained unigenes were annotated in the Gene Ontology (GO) database and Kyoto Encyclopedia of Genes and Genomes (KEGG) database for functional analysis. Differentially expressed genes (DEGs) were identified using R package edgeR, with significance cutoffs of log_2_(fold change) > 1 and *p* < 0.05. Pathway enrichment analysis of DEGs was performed using the KEGG database. The software TBtool (version 2.0) was used for further analysis of DEGs.

### 2.9. Gene Set Enrichment Analysis

Genes were sorted according to the degree of differential expression between the two sample groups. GSEA analysis (http://software.broadinstitute.org/gsea/index.jsp.) was conducted using the KEGG dataset, where FDR < 0.25, *p* < 0.05, and absolute normalized enrichment score (|NES|) > 1 indicate statistical significance.

### 2.10. Quantitative real-time PCR

Diatom cells from 50 mL culture were centrifuged and used for RNA extraction. The extracted RNA was subsequently reverse transcribed into cDNA using HiScript III RT SuperMix for qPCR (Vazyme Bio, Nanjing, China). The CFX96TM Real-Time System (Bio-Rad, Hercules, USA) and SYBR Green PCR reagents (Vazyme Bio, Nanjing, China) were used. The 18S ribosomal RNA housekeeping gene was selected as an internal control, and the relative gene transcription levels were determined using the 2^−ΔΔCt^ method [18]. Primers were designed using Primer3 (version 0.4.0) (http://frodo.wi.mit.edu/) and the sequences can be found in Table S1. Three biological replicates were included.

### 2.11. Metabolomic analysis

Microalgal cells were pelleted and resuspended in prechilled 80% methanol. Samples were incubated on ice for 30 s and then sonicated for 6 min. After centrifugation at 5,000 rpm, 4 °C for 1 min, the supernatant was freeze-dried and dissolved in 10% methanol. Finally, the solution was subjected to UHPLC-MS/MS analyses on a Vanquish UHPLC system (ThermoFisher, Germany) coupled with an Orbitrap Q Exactive HF mass spectrometer (ThermoFisher, Germany) at Novogene (Beijing, China). Raw files were processed using the Compound Discoverer 3.1 (CD3.1, ThermoFisher) to for peak alignment and quantitation. Metabolites were annotated using the KEGG (https://www.genome.jp/kegg/pathway.html), HMDB (https://hmdb.ca/metabolites), and LIPIDMaps (http://www.lipidmaps.org/) databases.

### 2.12. Statistical analysis

The experimental data in this study were obtained from three independent replicates (n = 3). One-way analysis of variance (ANOVA) was used for differential analysis (SPSS 25.0, Chicago). *p* < 0.05 was considered as statistical significance.

## 3. Results

### 3.1. Effects of IAA on diatom growth

Treatment of *P. tricornutum* with a wide range of IAA concentrations (100 pg L^-1^ to 100 μg L^-1^) for 24, 48, 72, and 96 h showed a growth promotion effect at 5 μg L^-1^ (Fig. 1). For instance, the growth promotion efficiency of 5 μg L^-1^ IAA was 10.6 % at 96 h, consistent to that of a previous study [19]. The growth promotion efficiency decreased at IAA higher than 5 μg L^-1^, with IAA of 100 μg L^-1^ showing an inhibitory effect. Therefore, 5 μg L^-1^ of IAA treatment was used in following experiments.

### 3.2. Effect of IAA on photosynthesis

We next assessed the effect of IAA on photosynthesis of *P. tricornutum*. Compared to the control group, the contents of chlorophyll *b* and carotenoids, but not chlorophyll *a*, were increased by 95% and 88%, respectively, after 24 h of IAA treatment, and by 1.46- and 2.48-fold, respectively, at 96 h (Fig. 2A, B and C). In addition, we observed a time-dependent increase in Fv/Fm, qP, and ETR in *P. tricornutum* cells supplemented with 5 μg L^-1^ of IAA compared to the control (Fig. 2D, E and F). At 96 h, Fv/Fm, qP, and ETR showed a 1.08-, 1.04-, and 1.05-fold increase compared to the control. By contrast, NPQ decreased with the time of treatment (Fig. 2G). Photosynthetic pigments are essential for photosynthesis and an increase in photosynthetic pigment content indicates an upregulation of photosynthetic pigment synthetase activity and capacity [20].

**Figure 1.**
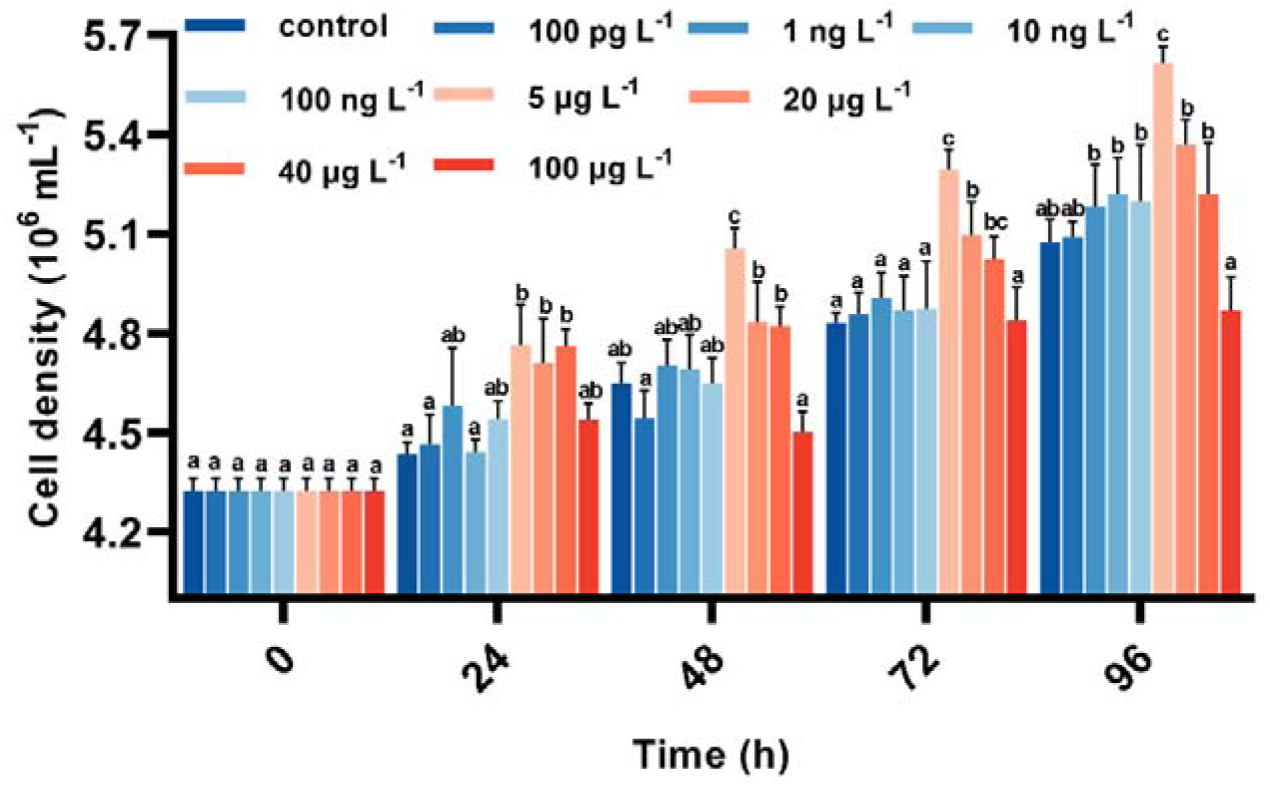
Effect of IAA on the growth rate of *P. tricornutum*. Error bars show the mean ± standard deviation of three replicates. Different letters indicate statistical significance (*p* < 0.05, Tukey’s test).

**Figure 2.**
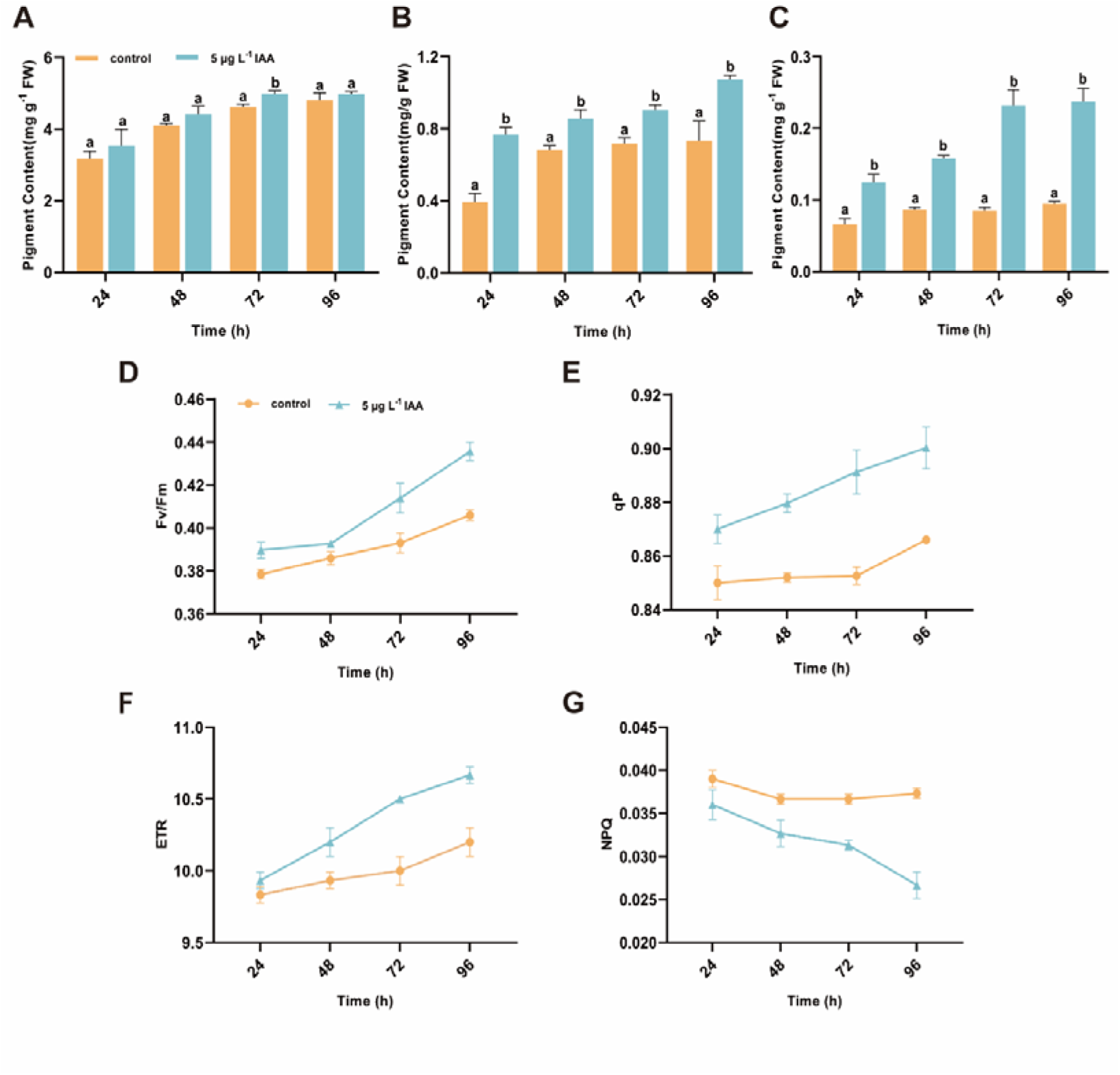
Effect of IAA on photosynthetic pigments and fluorescence. Chlorophyll *a* (A), chlorophyll *b* (B), carotenoids (C), Fv/Fm (D), qP (E), ETR (F), NPQ (G) were shown. Error bars show the mean ± standard deviation of three replicates. Different letters indicate statistical significance (*p* < 0.05, Tukey’s test).

### 3.3. Effects of IAA on respiratory oxygen consumption

To analyze the effect of IAA on respiration in *P. tricornutum*, we quantified the oxygen consumption under shading conditions. Consistent with the impact of IAA on photosynthetic rate, we observed an increase in net O_2_ release rate comparing the IAA-treated *P. tricornutum* to the control (Fig. 3A). In addition, the oxygen consumption rate of *P. tricornutum* in the control group was stable around 0.3 μmol·mg^-1^·min^-1^ during the experiment period. By contrast, IAA treatment led to a significant increase in oxygen consumption at all time points (Fig. 3B).

**Figure 3.**
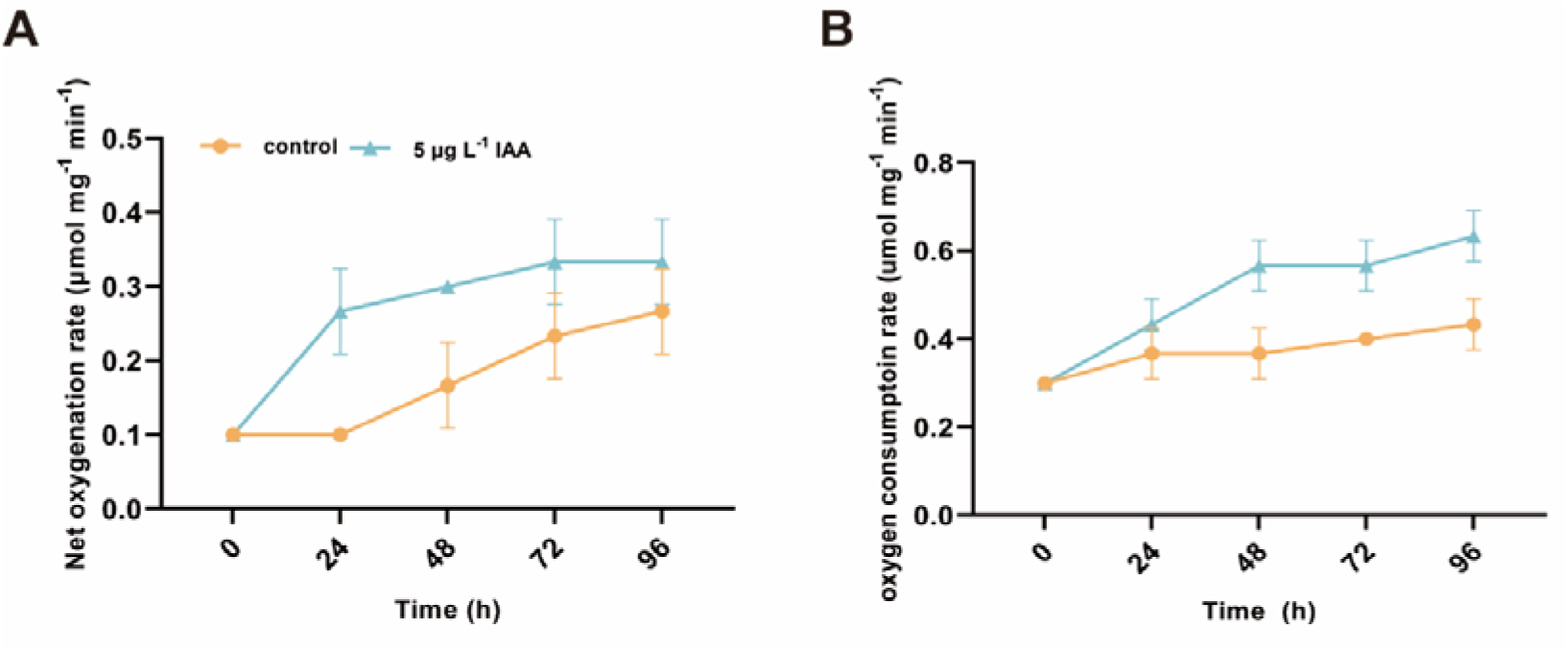
Effects of IAA on respiratory oxygen consumption. Changes of photosynthetic and respiration capacity index of *P. tricornutum* after 24, 48, 72, and 96 h exposure to IAA with net O_2_ release rate (A), respiratory O_2_ consumption rate (B).

### 3.4. Antioxidant enzymes activities of *P. tricornutum* after IAA treatment

To determine whether 5 μg L^-1^ of IAA causes stress responses in *P. tricornutum*, we examined the activity of antioxidant enzymes and the level of MDA. Compared to the control, IAA treatment led to a decrease in the activity of POD and CAT as well as the MDA content (Fig. 4). As these molecules are markers in oxidative stress responses, our data indicates that IAA does not introduce free radical stress in *P. tricornutum*.

**Figure 4.**
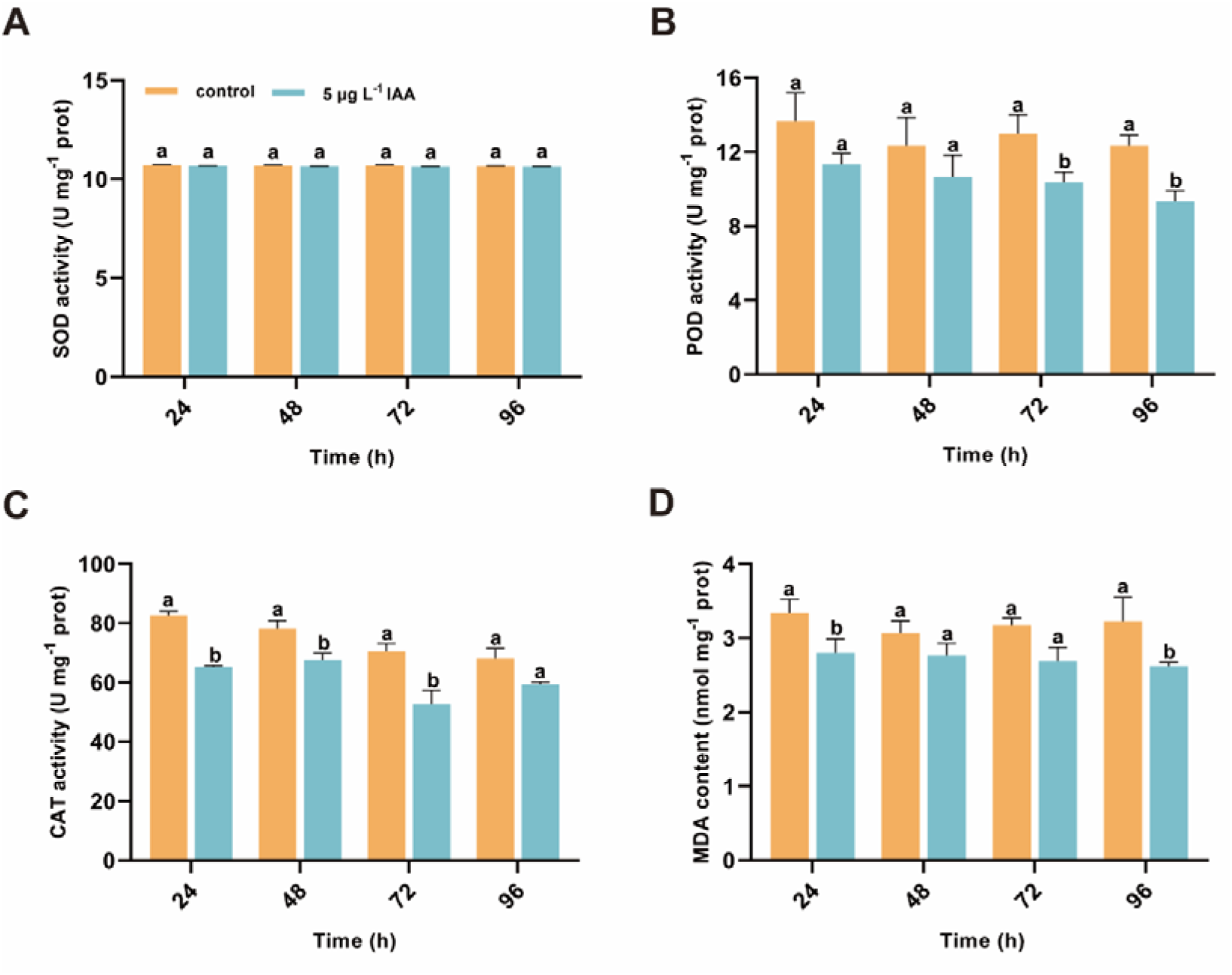
Antioxidant enzymes activities of *P. tricornutum* after IAA treatment. Effect of IAA on SOD activity (A), POD activity (B), CAT activity (C), and MDA content (D) in *P. tricornutum* cells. Error bars represent the standard error of the mean (n = 3). Different letters indicate statistical significance (*p* < 0.05, Tukey’s test).

### 3.5. Integrated transcriptomic and metabolomics analysis reveal the modulation of carbon metabolism by IAA

To elucidate the mechanism of the growth promotion effect of IAA, transcriptomics was carried out for *P. tricornutum* cells under 5 μg L^-1^ IAA treatment for 24 and 96 h, respectively (Fig. 5). In total, 1,085 and 1,076 genes were up- and down-regulated in *P. tricornutum* at 24 h, respectively (Fig. 5A). At 96 h, the numbers of differentially expressed genes (DEGs) were 586 (up-regulation) and 558 (down-regulation), respectively (Fig. 5B). A common set of 1,345 DEGs were found at both time points, including 25 genes coding for light-harvesting chlorophyll *a*/*b* binding proteins.

**Figure 5.**
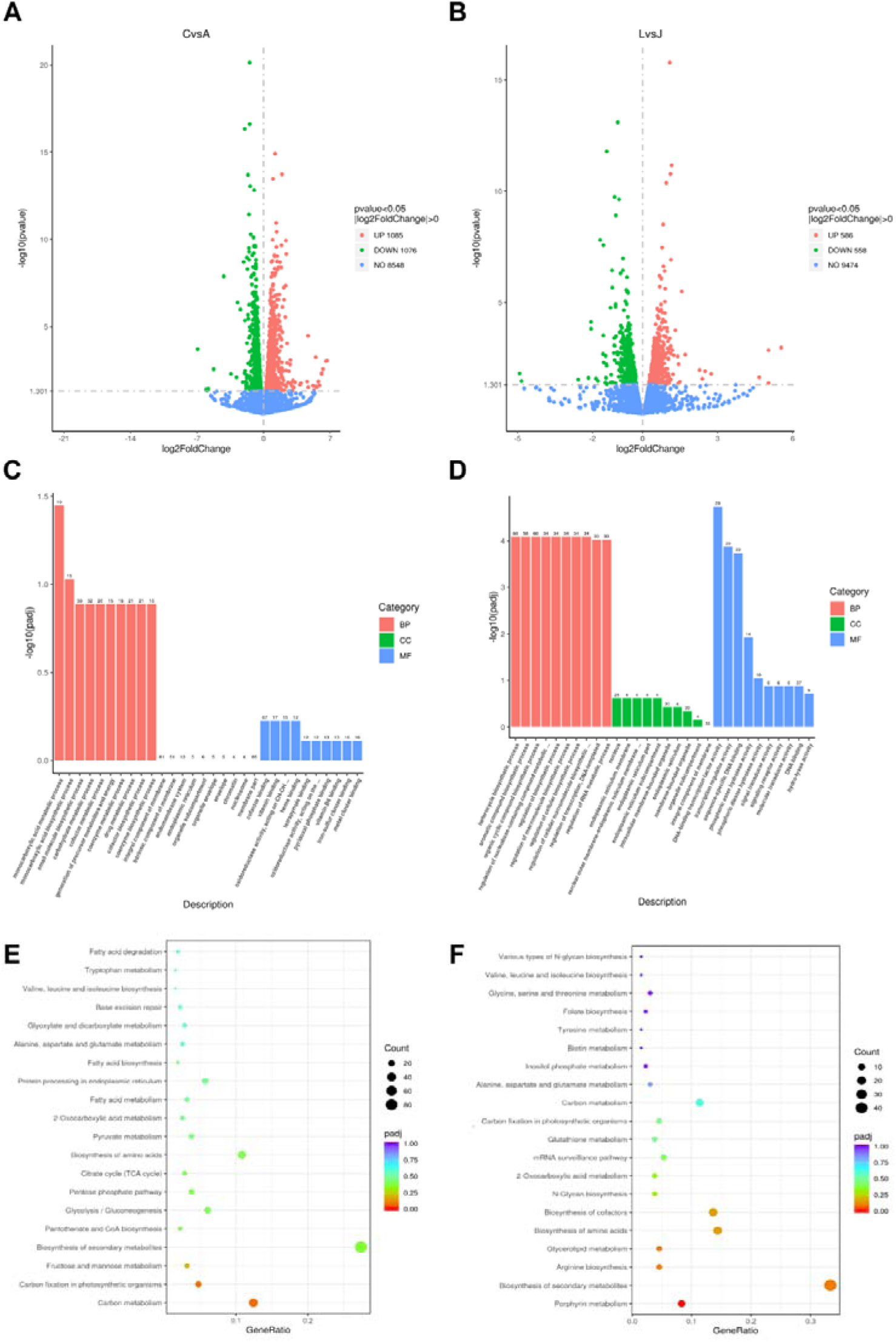
Analysis of differentially expressed genes (DEGs) in *P. tricornutum* under IAA treatment. A and B show the up-regulated (red) and down-regulated (green) genes at 24 and 96 h, respectively. C and D show the GO enrichment for 24 and 96 h, respectively. E and F show the KEGG enrichment of DEGs for 24 and 96 h, respectively.

GO analysis of DEGs resulted in a total of 30 significant terms (Fig. 5C and D). At 24 h, significant biological processes included monocarboxylic acid metabolic process (GO: 0032787), monocarboxylic acid biosynthetic process (GO: 0072330), small molecule biosynthetic process (GO: 0044283), and carbohydrate metabolic process (GO: 0005975). After 96 h of IAA treatment, heterocycle biosynthetic process (GO: 0018130), aromatic compound biosynthetic process (GO: 0019438), and organic cyclic compound biosynthetic process (GO:1901362) were significantly enriched. The differential enrichment patterns at the two time points indicate a time-dependent shift in cellular metabolism in the diatom cells, from carbon metabolism at 24 h to porphyrin metabolism (and other secondary metabolite synthesis, as well as arginine biosynthesis) at 96 h (Fig. 5).

We found multiple genes involved in photosynthetic carbon fixation are significantly upregulated at 24 h, revealing that IAA primarily affected the genes in the photosynthetic carbon fixation pathway of *P. tricornutum* at short time (24 h). Notably, this influence persisted for up to 96 h after IAA treatment, correlating with the enhanced photosynthetic efficiency, upregulation of photosynthetic pigments (e.g., porphyrin), and an increase in chlorophyll content (Fig. 2). Our enrichment of porphyrin metabolism at 96 h (Fig. 5) further supports the regulation of carbon metabolism, given the important role of porphyrins play in photosynthesis .

To reveal more deeply the major biochemical and signal transduction pathways involved under IAA treatment, we further performed metabolomics and integrated the data with transcriptome analysis. As expected, up-regulation of genes involved in carbon fixation by IAA was confirmed by the enhancement of corresponding metabolic pathways (Fig. 6A). For instance, we observed an up-regulation of PHATRDRAFT_50723 (δ-aminolevulinic acid dehydratase) gene and the accumulation of porphobilinogen (Fig. 6B), a precursor of chlorophyll.

**Figure 6.**
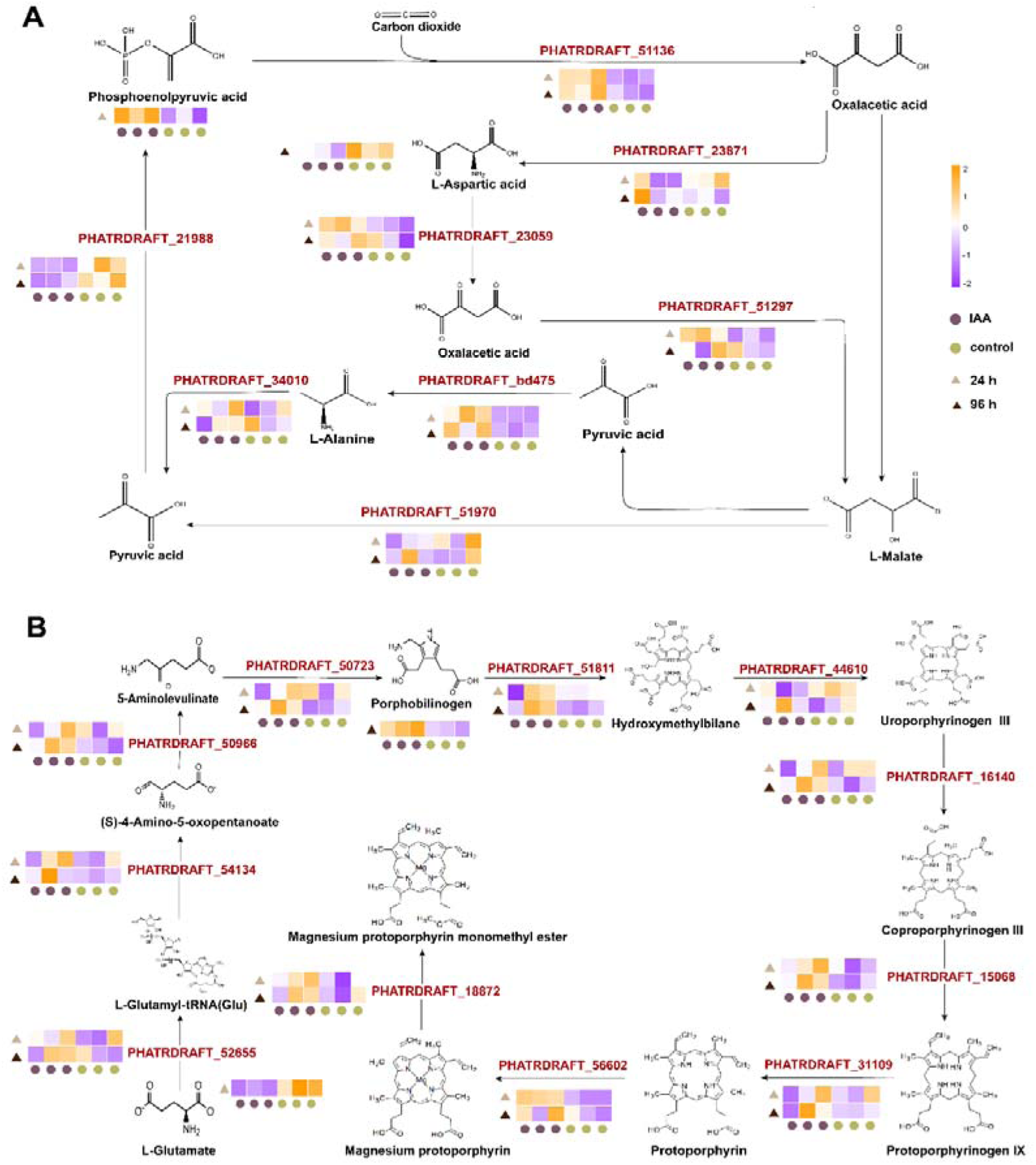
Heatmap of DEGs in the carbon sequestration pathway and porphyrin metabolism pathway. A and B show the DEGs in carbon sequestration pathway and porphyrin metabolism pathway, respectively.

We further performed gene set enrichment analysis (Fig. S1) and confirmed the differential enrichment patterns at the two time points. The prominent pathways identified at 24 h included inositol phosphate metabolism, pantothenate and CoA biosynthesis, autophagy, and the citrate cycle, while at 96 h, the key pathways encompassed oxocarboxylic acid metabolism, phenylalanine, tyrosine, and tryptophan biosynthesis, glycan biosynthesis, and porphyrin metabolism.

### 3.6. qPCR validation

We validated the transcriptomic results using qRT-PCR for four genes in carbon fixation and four in porphyrin metabolism (Fig. 7). Consistent with the transcriptome results, qRT-PCR showed up-regulation of genes by IAA in carbon fixation and porphyrin metabolism (Fig. 7). Of notes, most genes related to photosynthetic carbon fixation were up-regulated at both 24 and 96 h, with the exception of PHATRDRAFT_bd475 that was not up-regulated at 24 h and PHATRDRAFT_54279 that was not up-regulated at 96 h. In addition, the fours genes related to porphyrin metabolism were more significantly affected at 96 h. Interestingly, PHATRDRAFT_30690 was only significantly up-regulated at 96 h, and the other three genes were significantly updated at both time points.

**Figure 7.**
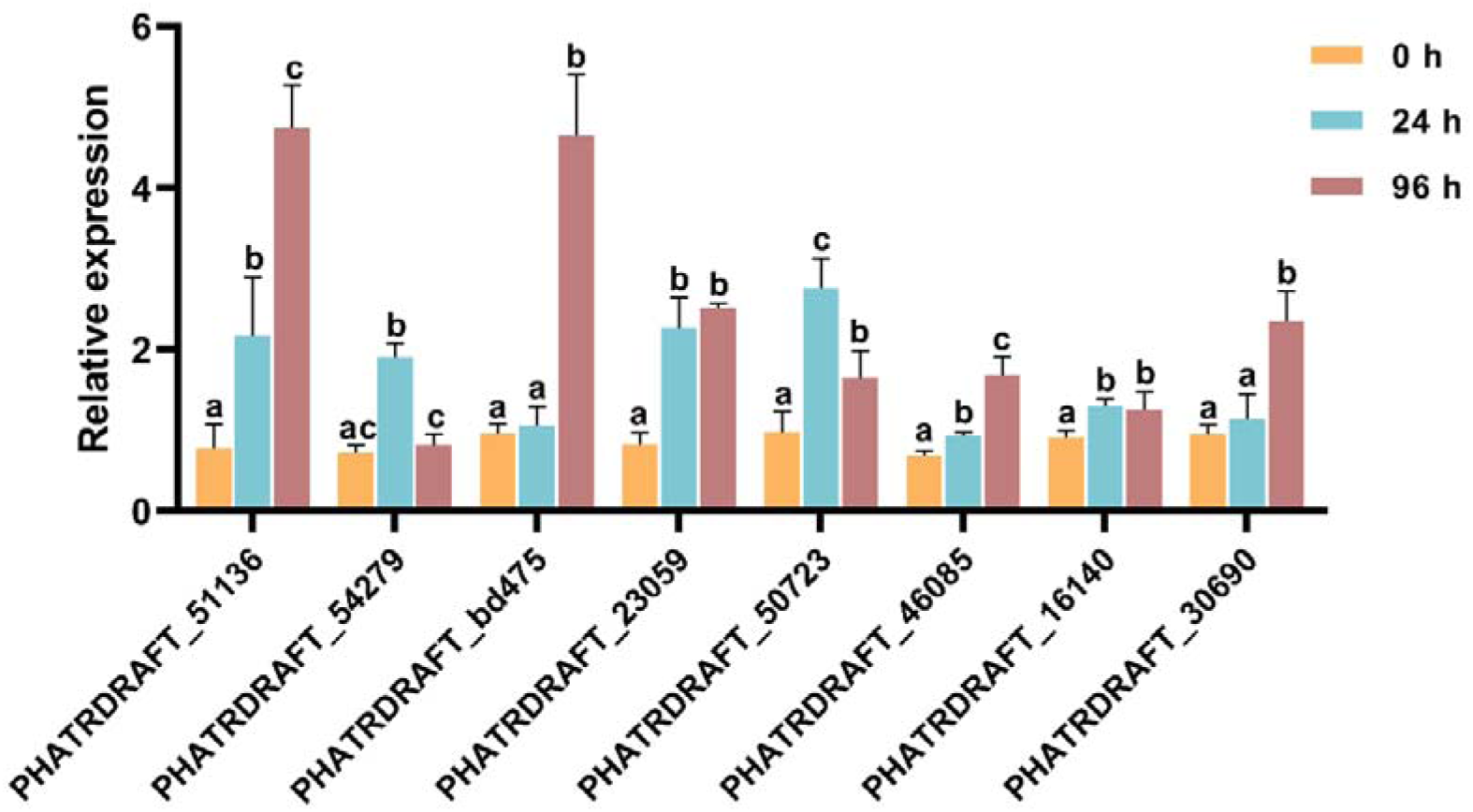
Real-time quantitative PCR analysis of genes involved in carbon fixation and terpenoid biosynthesis pathways. The first four genes are in carbon fixation pathway (count from left to right), and the last four are involved in terpenoid backbone biosynthesis (count from left to right). Data represented means ± SD for three independent replicates. Different letters indicated level of significance.

## 4. DISCUSSION

IAA has extensive applications in promoting the growth of microalgae [21]. In our experiment, we found that 5 μg L^-1^ IAA can strongly promote growth in *P. tricornutum*. However, high concentrations of IAA inhibit growth, as described in previous studies [19, 22]. Excess IAA could hinder cell division and differentiation [23] and induce the production of abscisic acid (ABA) and ethylene and thus inhibit the growth of algal cells [24].

We observed that IAA induce a considerable increase of chlorophyll and carotenoids in *P. tricornutum*, indicating an enhancement of photosynthesis and higher biomass production. Two mechanisms can explain the IAA-induced increase of photosynthetic pigment. First, IAA promotes the activation of genes responsible for chlorophyll and carotenoid biosynthesis [25, 26]. Second, IAA increases the efficiency of photosynthetic electron transport and energy conversion, ultimately leading to an increase in pigment synthesis [27]. In this mechanism, the pigment content reflects the efficiency of photosynthesis [28]. While chlorophyll *a* responds to the increased reactive oxygen species (ROS) in algae cells [29], chlorophyll *b* and carotenoids are particularly sensitive to IAA. In microalgae, IAA has been shown to increase the level of carotenoids [30], which serves as a defense mechanism to safeguard the photosynthetic system and maintain its efficiency [31].

Our data on chlorophyll fluorescence (e.g., an increase in Fv/Fm) also suggested that IAA promotes the photosynthetic capacity of *P. tricornutum* (Fig. 2). IAA could help to maintain a maximal quantum yield of PSII by improving the opening degree of the PSII reaction center and increasing the light utilization rate, as evidenced by the increase of ETR by IAA in our study. The light capture efficiency of PSII can be regulated by IAA via modulating the activity of light-activated enzymes in *P. tricornutum*. In addition, we saw a time-dependent gradual increase of qP (Fig. 2E), supporting the notion that IAA increases the proportion of open PSII reaction center and promotes PSII electron transfer capability (flow of electrons from the PSII oxidation side to the PSII reaction center). In addition, IAA enhances the self-protection ability of microalgae, as indicated by a decrease in NPQ (PSII regulated energy dissipation Y, an index of light protection) (Fig. 2G). Overall, we reason that 5 μg L^-1^ of IAA promotes photosynthetic capacity by increasing qP and decreasing NPQ. Efficient light energy capture is maintained with high photosynthetic pigment content, a high proportion of the open part of the PSII reaction center, and enhanced self-protection ability of PSII.

While IAA promotes photosynthesis, it also enhances the respiration of *P. tricornutum* (Fig. 3). As respiration is intimately linked to growth and cell division [32], the dark respiration rate can reflect growth rates [33]. Thus, the enhanced growth rate with 5 μg L^-1^ IAA treatment in this study (accelerated speed of cell division) results in an increase in respiratory rate.

In addition to photosynthesis and respiration, IAA had been shown to regulate the responses of microalgae and cyanobacteria under environmental stresses, including nutrient limitation, excessive salinity, heavy metals, and toxic compounds [34, 35]. We quantified the activities of antioxidant enzymes and found 5 μg L^-1^ of IAA did not impose stress on *P. tricornutum*. Thus, IAA activates stress tolerance mechanisms, enabling microorganisms to withstand and thrive in challenging environments [35].

Using transcriptomics, we found IAA upregulates a multitude of genes related to carbon metabolism and carbon fixation, similar to a prior study performed in *C. vulgaris* [36]. In accordance to an increased cell division, which requires a significant amount of energy, we also found upregulation of genes in energy metabolism pathways such as glycolysis, tricarboxylic acid (TCA) cycle, and pentose phosphate pathway. Glycolysis supplies energy for cellular activities and generates acetyl-CoA [36, 37], which enters the TCA cycle to produce NADH and FADH_2_ [38] or *de novo* fatty acid biosynthesis pathway in the plastid [39]. Our data on the upregulation of these pathways is consistent with the observation that IAA enhances the activity of enzymes in glycolysis and the TCA cycle [40]. Thus, IAA improves microalgal sugar biosynthesis and subsequent metabolism by regulating both gene expression and modulation of enzyme activities.

In addition to energy metabolic pathways, porphyrin metabolism is the most significantly enriched pathway at 96 h after IAA treatment. The transcriptome analysis identified 11 genes in the porphyrin metabolism (pti00860), including hemC, hemE and hemF, all of which were up-regulated. These genes encode enzymes that facilitate the conversion of uroporphyrinogen III to coproporphyrinogen III, protoporphyrinogen IX, and ultimately to protoporphyrin IX, serving as a crucial precursor for chlorophyll biosynthesis [41]. Consequently, the enhanced expression of these genes is expected to promote the synthesis of chlorophyll *b*. Moreover, earlier studies have also indicated that porphyrin and chlorophyll are vital antioxidant components in microalgae, serving to mitigate reactive oxygen species and alleviate oxidative stresses [16, 42]. This observation aligns with our measurements of antioxidant enzyme activities. Purines, such as adenine and guanine, have essential roles in the synthesis of ATPs, as well as serving as cell signaling regulators such as cAMP, cGMP, and NADH [43]. Several genes in purine metabolism were also upregulated. Purine metabolism is crucial for the growth and development of plants, as it provides the precursors for nucleic acids, phospholipids, polysaccharides and secondary metabolites [44, 45]. This again supports the role of IAA in promoting growth.

Our findings also revealed an enhancement in the biosynthesis of amino acids, particularly arginine, at 96 h. Enrichment of multiple amino acid synthesis pathways was also found in GSEA analysis (Fig. S1). IAA can induce the expression of genes and the activity of enzymes in amino acid metabolism [46]. For instance, IAA resulted in increased levels of most amino acids, with the exception of methionine, in in *Chlorella* [47]. Among the amino acids, arginine plays a major role in combating oxidative stres [48–50], which is consistent with our results.

Overall, IAA stimulated the yields of microalgal biomass by enhancing carbon fixation and porphyrin metabolism as well as generating more energy by upregulating energy metabolisms. Future investigations should prioritize large-scale trials to evaluate the promise of adding IAA into outdoor algal ponds.

## 5. CONCLUSION

We found the optimal IAA concentration for enhancing the growth rate of *P. tricornutum* was 5 μg L^-1^. IAA treatment led to significant changes in photosynthetic pigments, fluorescence parameters, and photosynthetic oxygen evolution. In addition, transcriptomics revealed that IAA promotes the growth of *P. tricornutum* by regulating genes involved in the photosynthetic carbon fixation system and porphyrin metabolism.

## ACKNOWLEDGMENTS

This work was supported by Zhejiang Provincial Natural Science Foundation of China (LY23D060001), the National Key R&D Program of China (2018YFD0901504), and the National Natural Science Foundation of China (31670402). We also thank Hai-Hong Wen and Zhi-Wei Hu (National and Local Joint Engineering Research Center of Ecological Treatment Technology for Urban Water Pollution, Wenzhou University) for assistance with the experiments.

## CREDIT AUTHORSHIP CONTRIBUTION STATEMENT

Yu-Ting Chen: Investigation, Writing – original draft. Dong-Sheng Zhao: Investigation, Software. Xiao-Li Liu: Investigation. Huan-Yang: Investigation. Run- Ze Gu: Data curation, Validation. Na Li: Resources. Xiufeng Yan: Supervision. Hui- Xi Zou: Funding acquisition, Writing – review & editing, Supervision.

## DATA AVAILABILITY

Data will be made available on request.

## REFERENCES

[1] S. Malviya, E. Scalco, S. Audic, F. Vincent, A. Veluchamy, J. Poulain, P. Wincker, D. Iudicone, C. de Vargas, L. Bittner, A. Zingone, C. Bowler, Insights into global diatom distribution and diversity in the world’s ocean, Proceedings of the National Academy of Sciences of the United States of America, 113 (2016) E1516–1525.

[2] S.-H. Lee, R. Karawita, M. Affan, J.-B. Lee, K.-W. Lee, J.-S. Kim, D.-W. Kim, Y.-J. Jeon, Potential of Benthic Diatoms Achnanthes longipes, Amphora coffeaeformisand Navicula sp. (Bacillariophyceae) as Antioxidant Sources, Algae, 24 (2009) 47–55.

[3] P. Chelf, Environmental control of lipid and biomass production in two diatom species, Journal of Applied Phycology, 2 (1990) 121–129.

[4] B. Aslanbay Guler, I. Deniz, Z. Demirel, S.S. Oncel, E. Imamoglu, Comparison of different photobioreactor configurations and empirical computational fluid dynamics simulation for fucoxanthin production, Algal Research, 37 (2019) 195–204.

[5] A.J. Haagen-Smit, W.D. Leech, W.R. Bergen, Estimation, Isolation and Identification of Auxins in Plant Material, Science, 93 (1941) 624–625.

[6] S. Abel, A. Theologis, Odyssey of Auxin, Cold Spring Harbor Perspectives in Biology, 2 (2010) a004572.

[7] A. Poulet, V. Kriechbaumer, Bioinformatics Analysis of Phylogeny and Transcription of TAA/YUC Auxin Biosynthetic Genes, International Journal of Molecular Sciences, 18 (2017) 1791.

[8] F. Romani, Origin of TAA Genes in Charophytes: New Insights into the Controversy over the Origin of Auxin Biosynthesis, Frontiers in Plant Science, 8 (2017).

[9] R.W. Hunt, S. Chinnasamy, K.C. Das, The Effect of Naphthalene-Acetic Acid on Biomass Productivity and Chlorophyll Content of Green Algae, Coccolithophore, Diatom, and Cyanobacterium Cultures, Applied Biochemistry and Biotechnology, 164 (2011) 1350–1365.

[10] G.-H. Dao, G.-X. Wu, X.-X. Wang, T.-Y. Zhang, X.-M. Zhan, H.-Y. Hu, Enhanced microalgae growth through stimulated secretion of indole acetic acid by symbiotic bacteria, Algal Research, 33 (2018) 345–351.

[11] J.J. Tate, M.T. Gutierrez-Wing, K.A. Rusch, M.G. Benton, The Effects of Plant Growth Substances and Mixed Cultures on Growth and Metabolite Production of Green Algae Chlorella sp.: A Review, Journal of Plant Growth Regulation, 32 (2013) 417–428.

[12] C. Yi, R.T.-H. Skye, M.S. Peer, Phaeodactylum tricornutum microalgae as a rich source of omega-3 oil: Progress in lipid induction techniques towards industry adoption, Food Chemistry, 297 (2019) 124937.

[13] L. Gouveia, A.C. Oliveira, Microalgae as a raw material for biofuels production, Journal of Industrial Microbiology and Biotechnology, 36 (2009) 269–274.

[14] Y. Wei, N. Zhu, M. Lavoie, J. Wang, H. Qian, Z. Fu, Copper toxicity to Phaeodactylum tricornutum: a survey of the sensitivity of various toxicity endpoints at the physiological, biochemical, molecular and structural levels, Biometals, 27 (2014) 527–537.

[15] S. Zuo, S. Zhou, L. Ye, S. Ma, Synergistic and antagonistic interactions among five allelochemicals with antialgal effects on bloom-forming Microcystis aeruginosa, Ecological Engineering, 97 (2016) 486–492.

[16] J.-Q. Xiong, M.B. Kurade, R.A.I. Abou-Shanab, M.-K. Ji, J. Choi, J.O. Kim, B.-H. Jeon, Biodegradation of carbamazepine using freshwater microalgae Chlamydomonas mexicana and Scenedesmus obliquus and the determination of its metabolic fate, Bioresource Technology, 205 (2016) 183–190.

[17] X. Wang, J. Miao, L. Pan, Y. Li, Y. Lin, J. Wu, Toxicity effects of p- choroaniline on the growth, photosynthesis, respiration capacity and antioxidant enzyme activities of a diatom, Phaeodactylum tricornutum, Ecotoxicology and Environmental Safety, 169 (2019) 654–661.

[18] K.J. Livak, T.D. Schmittgen, Analysis of relative gene expression data using real-time quantitative PCR and the 2(-Delta Delta C(T)) Method, Methods (San Diego, Calif.), 25 (2001) 402–408.

[19] Z. Wang, J. Mou, Z. Qin, Y. He, Z. Sun, X. Wang, C.S.K. Lin, An auxin-like supermolecule to simultaneously enhance growth and cumulative eicosapentaenoic acid production in Phaeodactylum tricornutum, Bioresource Technology, 345 (2022) 126564.

[20] N. Baker, Chlorophyll Fluorescence: A Probe of Photosynthesis In Vivo, Annual Review of Plant Biology, 59 (2008) 89–113.

[21] Z. Yu, M. Song, H. Pei, L. Jiang, Q. Hou, C. Nie, L. Zhang, The effects of combined agricultural phytohormones on the growth, carbon partitioning and cell morphology of two screened algae, Bioresource Technology, 239 (2017) 87–96.

[22] A. Gonzalez-Garcinuno, J.M. Sanchez-Alvarez, M.A. Galan, E.M. Martin del Valle, Understanding and optimizing the addition of phytohormones in the culture of microalgae for lipid production, Biotechnology Progress, 32 (2016) 1203–1211.

[23] W.H.A. Hakim, T. Erfianti, A.N. Dhiaurahman, K.Q. Maghfiroh, R. Amelia, I. Nurafifah, D. Kurniantob, D.U. Siswanti, E.A. Suyono, S. Marno, I. Devi, The Effect of IAA Phytohormone (Indole-3- Acetic Acid) on the Growth, Lipid, Protein, Carbohydrate, and Pigment Content in Euglena sp, Malaysian Journal of Fundamental and Applied Sciences, 19 (2023) 513–524.

[24] A. Udayan, M. Arumugam, Selective enrichment of Eicosapentaenoic acid (20:5n-3) in N. oceanica CASA CC201 by natural auxin supplementation, Bioresource Technology, 242 (2017) 329–333.

[25] R. Defez, A. Andreozzi, S. Romano, G. Pocsfalvi, I. Fiume, R. Esposito, C. Angelini, C. Bianco, Bacterial IAA-Delivery into Medicago Root Nodules Triggers a Balanced Stimulation of C and N Metabolism Leading to a Biomass Increase, Microorganisms, 7 (2019) 403.

[26] M. Ruiz-Sola, M. Rodríguez Concepción, Carotenoid biosynthesis in Arabidopsis: a colorful pathway, The arabidopsis book, 10 (2012) e0158.

[27] B.S. Kumudini, S.V. Patil, Role of Plant Hormones in Improving Photosynthesis, Photosynthesis, Productivity and Environmental Stress, (2019) 215–240.

[28] R. Santos, Effects of Cadmium on Growth, Photosynthetic Pigments, Photosynthetic Performance, Biochemical Parameters and Structure of Chloroplasts in the Agarophyte Gracilaria domingensis (Rhodophyta, Gracilariales), American Journal of Plant Sciences, 03 (2012) 1077–1084.

[29] Y. Xiao, X. Jiang, Y. Liao, W. Zhao, P. Zhao, M. Li, Adverse physiological and molecular level effects of polystyrene microplastics on freshwater microalgae, Chemosphere, 255 (2020) 126914.

[30] R. Czerpak, A. Bajguz, Stimulatory Effect of Auxins and Cytokinins on Carotenes, with Differential Effects on Xanthophylls in the Green Alga Chlorella Pyrenoidosa Chick, Acta Societatis Botanicorum Poloniae, 66 (1997) 41–46.

[31] D.B. Rodrigues, A.Z. Mercadante, L.R.B. Mariutti, Marigold carotenoids: Much more than lutein esters, Food Research International, 119 (2019) 653–664.

[32] O. Perez-Garcia, F.M.E. Escalante, L.E. de-Bashan, Y. Bashan, Heterotrophic cultures of microalgae: Metabolism and potential products, Water Research, 45 (2011) 11–36.

[33] R. Geider, B. Osborne, Respiration and microalgal growth: a review of the quantitative relationship between dark respiration and growth, New Phytologist, 112 (2006) 327–341.

[34] C.-Y. Tan, I.C. Dodd, J.E. Chen, S.-M. Phang, C.F. Chin, Y.-Y. Yow, S. Ratnayeke, Regulation of algal and cyanobacterial auxin production, physiology, and application in agriculture: an overview, Journal of Applied Phycology, 33 (2021) 2995–3023.

[35] M.R. Gauthier, G.N.A. Senhorinho, J.A. Scott, Microalgae under environmental stress as a source of antioxidants, Algal Research, 52 (2020) 102104.

[36] J.-M. Xu, J.-Q. Xiong, Boosting the yields of microalgal biomass and high-value added products by phytohormones: A mechanistic insight using transcriptomics, Journal of Cleaner Production, 393 (2023) 136175.

[37] H. Li, J. Tan, Y. Mu, J. Gao, Lipid accumulation of Chlorella sp. TLD6B from the Taklimakan Desert under salt stress, PeerJ, 9 (2021) e11525.

[38] L. Alipanah, J. Rohloff, P. Winge, A.M. Bones, T. Brembu, Whole-cell response to nitrogen deprivation in the diatom Phaeodactylum tricornutum, Journal of Experimental Botany, 66 (2015) 6281–6296.

[39] G. Müller, S. Wied, S. Over, W. Frick, Inhibition of lipolysis by palmitate, H2O2 and the sulfonylurea drug, glimepiride, in rat adipocytes depends on cAMP degradation by lipid droplets, Biochemistry, 47 (2008) 1259–1273.

[40] R. Casanova-Sáez, E. Mateo-Bonmatí, K. Ljung, Auxin Metabolism in Plants, Cold Spring Harbor Perspectives in Biology, 13 (2021) a039867.

[41] X. Xiao, W. Li, S. Li, X. Zuo, J. Liu, L. Guo, X. Lu, L. Zhang, The Growth Inhibition of Polyethylene Nanoplastics on the Bait-Microalgae Isochrysis galbana Based on the Transcriptome Analysis, Microorganisms, 11 (2023) 1108.

[42] S. Thunell, Porphyrins, porphyrin metabolism and porphyrias. I. Update, Scandinavian Journal of Clinical and Laboratory Investigation, 60 (2000) 509–540.

[43] J. Mo, Z. Ma, S. Yan, N.K.M. Cheung, F. Yang, X. Yao, J. Guo, Metabolomic profiles in a green alga (Raphidocelis subcapitata) following erythromycin treatment: ABC transporters and energy metabolism, Journal of Environmental Sciences, 124 (2023) 591–601.

[44] J.-J. Park, H. Wang, M. Gargouri, R.R. Deshpande, J.N. Skepper, F.O. Holguin, M.T. Juergens, Y. Shachar-Hill, L.M. Hicks, D.R. Gang, The response of Chlamydomonas reinhardtii to nitrogen deprivation: a systems biology analysis, The Plant Journal, 81 (2015) 611–624.

[45] C. Stasolla, R. Katahira, T.A. Thorpe, H. Ashihara, Purine and pyrimidine nucleotide metabolism in higher plants, Journal of Plant Physiology, 160 (2003) 1271–1295.

[46] J. Kim, K. Harter, A. Theologis, Protein–protein interactions among the Aux/IAACproteins, PNAS, 94 (1997) 11786–11791.

[47] W.A. Fathy, H. AbdElgawad, A.H. Hashem, E. Essawy, E. Tawfik, A.A. Al-Askar, M.S. Abdelhameed, O. Hammouda, K.N.M. Elsayed, Exploring Exogenous Indole-3-acetic Acid’s Effect on the Growth and Biochemical Profiles of Synechocystis sp. PAK13 and Chlorella variabilis, Molecules, 28 (2023) 5501.

[48] M. Liang, Z. Wang, H. Li, L. Cai, J. Pan, H. He, Q. Wu, Y. Tang, J. Ma, L. Yang, l-Arginine induces antioxidant response to prevent oxidative stress via stimulation of glutathione synthesis and activation of Nrf2 pathway, Food and Chemical Toxicology, 115 (2018) 315–328.

[49] N.M. Silveira, R.V. Ribeiro, S.F.N. de Morais, S.C.R. de Souza, S.F. da Silva, A.B. Seabra, J.T. Hancock, E.C. Machado, Leaf arginine spraying improves leaf gas exchange under water deficit and root antioxidant responses during the recovery period, Plant Physiology and Biochemistry, 162 (2021) 315–326.

[50] J. Fuhrmann, V. Subramanian, Douglas J. Kojetin, Paul R. Thompson, Activity-Based Profiling Reveals a Regulatory Link between Oxidative Stress and Protein Arginine Phosphorylation, Cell Chemical Biology, 23 (2016) 967–977.

