## Supplemental Figure 1, Supplemental Table 1 for "Physiological, transcriptomic and metabolomic responses of the marine diatom *Phaeodactylum tricornutum* to auxin"

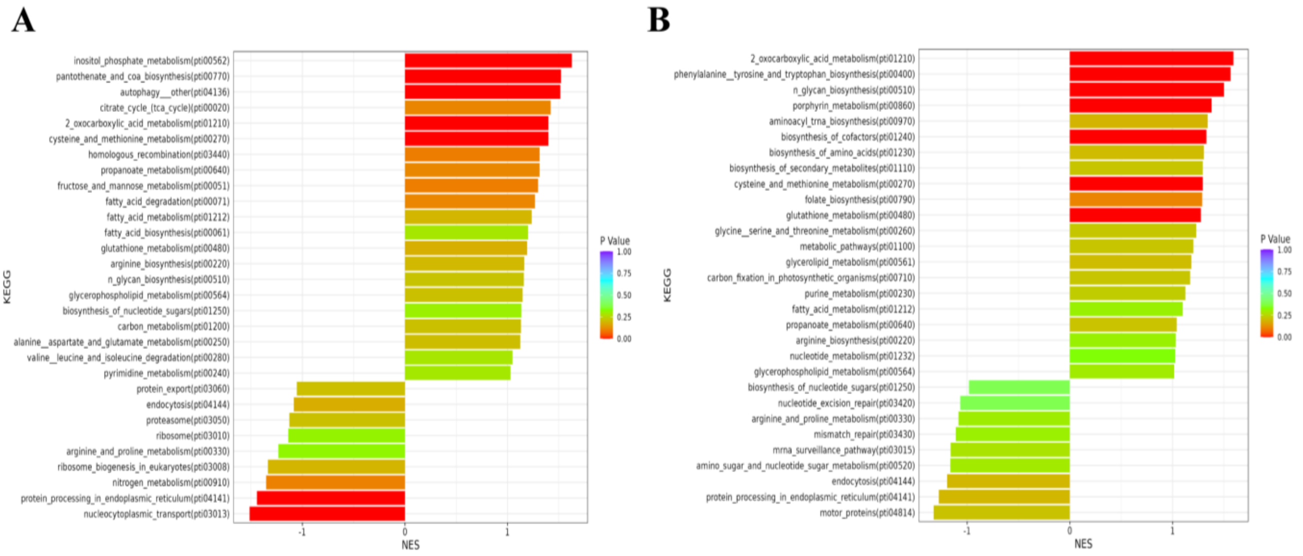


**Figure S1**. **Analysis of differentially expressed genes (DEGs) in *P. tricornutum* under IAA treatment.** A and B show the GSEA results for 24 h and 96 h, respectively.

**Table S1**. Primers used in this study.

| Primer names | 5’ → 3’ |
|---|---|
| PHATRDRAFT_51136_for | CTTCTCAAGCTTAGCCCCG |
| PHATRDRAFT_51136_rev | GAAGACAACCGGACACTCTG |
| PHATRDRAFT_54279_for | CCGTCTTTGCCACTTTTCAG |
| PHATRDRAFT_54279_rev | ATGCATTTCGGAGACAGGAG |
| PHATRDRAFT_bd475_for | CCAATTCACAAGGTTTAGCCG |
| PHATRDRAFT_bd475_rev | ATCGGGTATTGAGGAATTGGG |
| PHATRDRAFT_23059_for | TTACATTCCCACACCCACG |
| PHATRDRAFT_23059_rev | AGCAATCCGTCCAAATCCAG |
| PHATRDRAFT_50723_for | AAAGGTGTGGATGCCGATAG |
| PHATRDRAFT_50723_rev | AGGGATAAATGAAGTTGCTGGG |
| PHATRDRAFT_46085_for | CTCCTACATTTCACTACCCTGG |
| PHATRDRAFT_46085_rev | TTCCGAGTCCTGCAATTGAG |
| PHATRDRAFT_16140_for | GGAATATCAGGACTACAAGGCG |
| PHATRDRAFT_16140_rev | AAAATAGCCGCATCCAAATCG |
| PHATRDRAFT_30690_for | GAATTGTTTGAATGCGGGTAGAG |
| PHATRDRAFT_30690_rev | CAATGGTCCAGGTCATATCGG |
